# Curiosity and mesolimbic functional connectivity drive information seeking in real life

**DOI:** 10.1101/2022.01.28.478038

**Authors:** Kathrin C. J. Eschmann, Duarte F. M. M. Pereira, Ashvanti Valji, Vera Dehmelt, Matthias J. Gruber

**Author notes:** equal contribution (Kathrin C. J. Eschmann, Duarte F. M. M. Pereira).

## Abstract

Curiosity reflects the intrinsic motivation of an individual to seek information in order to close information gaps. Laboratory-based experiments have shown that both curiosity and information seeking are associated with enhanced neural dynamics in the mesolimbic dopaminergic circuit. However, it is unclear whether curiosity and its associated neural dynamics in the dopaminergic circuit drive information seeking in real life. The present study investigated (i) whether curiosity traits predict different characteristics of real-life information seeking and (ii) whether functional connectivity within the mesolimbic dopaminergic circuit is associated with information seeking outside of the laboratory. Up to 15 month before the COVID-19 pandemic, we conducted curiosity and anxiety questionnaires as well as a 10-minute resting-state fMRI session. In a follow-up survey early during the COVID-19 pandemic, participants repeated the questionnaires and filled out an additional questionnaire about their COVID-19-related information seeking. Curiosity but not anxiety remained stable over time. Individual differences in curiosity were positively associated with the frequency of information-seeking behaviour. Anxiety during the pandemic was not linked to any characteristics of real-life information seeking. Interestingly, the frequency of information seeking was also independently predicted by individual differences in resting-state functional connectivity between the ventral tegmental area and the nucleus accumbens. The present translational study paves the way for future studies on the role of curiosity in real-life information seeking by showing that curiosity drives information seeking in real-life situations and that the curiosity-promoting mesolimbic dopaminergic functional network supports the frequency of real-life information-seeking behaviour.

**SIGNIFICANCE STATEMENT:** Curiosity is a key driver of learning and information seeking in everyday life. However, the temporal stability of curiosity traits, their relationship to real-life information seeking, and the associated dopaminergic brain activity are poorly understood. The present study provides evidence that curiosity traits are stable over time – even through a major event, such as the COVID-19 pandemic – and that both curiosity and intrinsic functional connectivity within the mesolimbic dopaminergic circuit are associated with the frequency of real-life information seeking. These findings contribute to a better understanding of cognitive and neural differences that shape how individuals seek out information and may offer the opportunity to help individuals with suboptimal information-seeking behaviour that negatively affects their well-being or mental health.

## INTRODUCTION

Curiosity - the desire to learn about specific information without any apparent extrinsic rewards - is a driver of information seeking that has been studied in a variety of laboratory settings (1–3). It has been proposed that curiosity can be elicited by the detection of information gaps or so-called information prediction errors (4, 5). Subsequent information seeking serves to close these information gaps and reduce uncertainty (1, 4, 6). Notably, information gaps might result in anxiety instead of curiosity if the current state of uncertainty is perceived as a threat or the necessary resources for successful uncertainty resolution are missing (4, 7, 8). Although laboratory-based research shows that curiosity is one of the key drivers of information seeking, it is unknown whether these findings translate to information seeking in everyday life where genuine knowledge acquisition takes place. An initial hint comes from a study showing that deprivation sensitivity, that is, a subtype of curiosity reflecting the tendency to seek information in order to close information gaps, was associated with the creation of knowledge networks during exploration of Wikipedia articles (9). Given that participants were able to explore individually chosen topics that they were interested in, it remains an open question whether curiosity drives real-life information seeking for a specific topic that is novel and personally relevant across all information seekers.

Curiosity about various types of information, such as trivia answers, magic tricks, and morbid images, has been associated with activation in the mesolimbic dopaminergic circuit (10–15). For instance, high ratings of curiosity were shown to be accompanied by increased activity in the ventral tegmental area (VTA) and the nucleus accumbens (NAcc; 10), supporting the idea that the dopaminergic system supports the drive to seek information (16). Consistent with this interpretation, information seeking itself has also been related to increases in activity and functional connectivity within the mesolimbic dopaminergic system (14, 17). Specifically, functional connectivity between the VTA and NAcc increased during presentation of a cue that informs about upcoming information, indicating that these dopaminergic brain regions form a network that is important for information-seeking decisions (17). Furthermore, mesolimbic dopaminergic brain areas show task-independent intrinsic functional connectivity amongst each other and its strength varies between individuals (18). Based on these findings, the question arises whether intrinsic functional connectivity between key regions of the mesolimbic dopaminergic circuit, such as the VTA and NAcc, is also associated with curiosity-driven information seeking in real life.

The recent COVID-19 pandemic offered a unique opportunity to study curiosity, the underlying neural dynamics in the mesolimbic dopaminergic system, and their relationship to information seeking in real life. The novelty of the SARS-CoV-2 virus and the uncertainty about personally relevant health, social, and economic consequences introduced information gaps about COVID-19-related information to the world-wide population. Initial research has suggested that states of curiosity drive information seeking about COVID-19-related news by maximising personal utility as shown in an online experiment, in which participants’ curiosity was measured as the willingness to wait for COVID-19-related information (19). However, it remains an open question whether individual differences related to how strongly people generally express curiosity, that is curiosity traits, predict information seeking of COVID-19-related news. It is also unclear whether curiosity traits, which are thought to be rather stable personality characteristics, are unaffected by major disruptions to daily life, such as during the beginning of the COVID-19 pandemic. Based on the overarching influence of the early COVID-19 pandemic on physiological and psychological well-being (20, 21), the temporal stability of curiosity traits should be especially probed under these life-changing circumstances that might be perceived as threatening and consequently enhance anxiety levels. Thus far, anxiety levels during the COVID-19 pandemic have been associated with a reduction in the willingness to wait for information (19) but also with increased information-seeking behaviour (22, 23). Anxiety predicted COVID-19-related information seeking (22) but was not related to information seeking when obsessive compulsive behaviour was taken into account (23), indicating that anxiety is an important control variable when determining the key drivers of real-life information seeking. Altogether, it is not clear whether individual differences in curiosity solely or jointly with anxiety affect individual differences in real-life information seeking about COVID-19-related news.

In the present study, participants of a previous neuroimaging study were re-invited to participate in a follow-up survey during the beginning of the COVID-19 pandemic (Figure 1). Up to 15 months before the introduction of COVID-19 lockdown restrictions, individual curiosity and anxiety scores were measured with the help of questionnaires (24, 25) and a 10-minute restingstate functional magnetic resonance imaging (fMRI) session was conducted. After the introduction of COVID-19 lockdown restrictions, participants repeated the curiosity and anxiety questionnaires and filled out an additional questionnaire about the characteristics of their COVID-19-related information-seeking behaviour. More specifically, the information seeking questionnaire assessed the frequency, detail, duration, and diversity of information seeking during the first month of lockdown (Figure 1). We predicted that curiosity measurements should not change from before to during the COVID-19 pandemic if curiosity traits measured with the Five-Dimensional Curiosity Scale (24) reflect temporally stable personality traits. In contrast, anxiety levels may change during the COVID-19 pandemic. Given that curiosity was shown to be a key driver of information seeking in laboratory-based experiments (1–3), we hypothesised that individual differences in curiosity may predict characteristics of pandemic-related information seeking in real life independently of anxiety levels. Furthermore, characteristics of information seeking should be positively associated with intrinsic functional connectivity between the VTA and NAcc, which previously has been related to curiosity and information seeking in experimental settings (10, 17).

**Figure 1.**
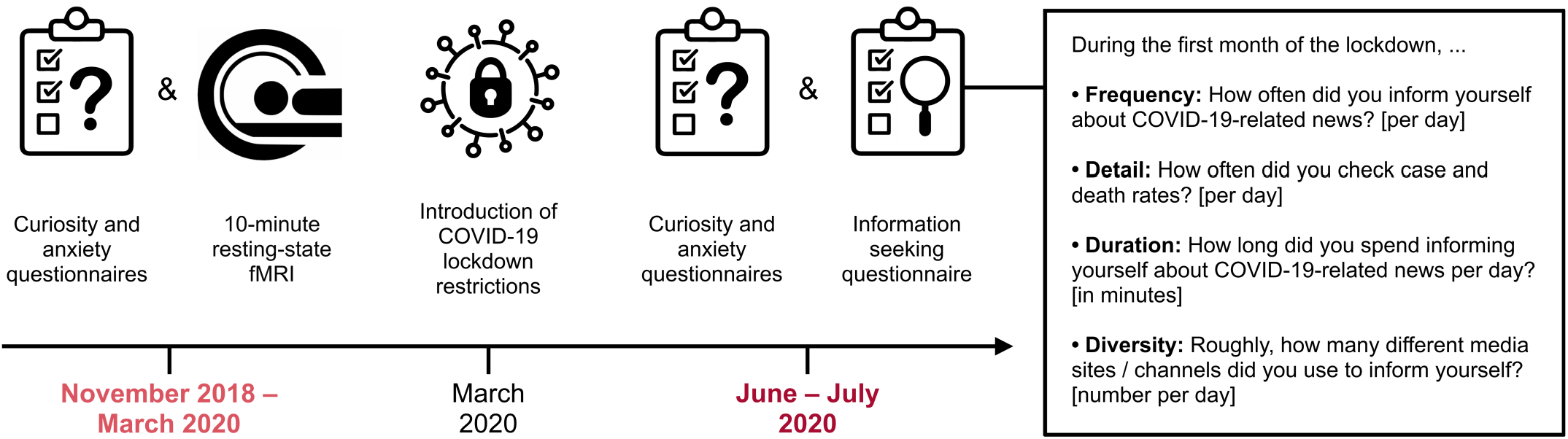
Overview of the experimental timeline and details of the COVID-19-related information seeking questionnaire. Curiosity and anxiety questionnaires as well as 10-minute resting-state functional magnetic resonance imaging (fMRI) were conducted up to 15 months before COVID-19 lockdown restrictions were introduced in the United Kingdom. During the COVID-19 pandemic, participants repeated the questionnaires and filled out an additional questionnaire about their information-seeking behaviour during the first month of lockdown. Labels for each item of the information seeking questionnaire were not presented to participants.

## RESULTS

### Are curiosity and anxiety stable over time?

Curiosity as measured with the Five-dimensional Curiosity Scale (24) did not change from before to during the - COVID-19 pandemic (*t*(28) = 0.13, *p* = .895, *d* = 0.02, two-tailed), indicating that curiosity traits remained stable over time (Figure 2A). This was further supported by an intraclass correlation coefficient (ICC) that indicated good temporal stability of the five-dimensional curiosity scores (ICC = 0.84, 95% CI [.70, .91]). Notably, participants also showed no difference from before to during the pandemic in any of the five-dimensional curiosity subscales (all FDR-adjusted *p*-values > .534, two-tailed) as revealed by ICC values indicating moderate to good stability for joyous exploration (ICC = 0.81, 95% CI [.65, | .90]), deprivation sensitivity (ICC = 0.67, 95% CI [.39,.83]), stress tolerance (ICC = 0.81, 95% CI [.65, .90]), social curiosity (ICC = 0.84, 95% CI [.69, .91]), and thrill-seeking (ICC = 0.83, 95% CI [.69, .91]). In contrast, anxiety as measured by the short form of the State-Trait Anxiety Inventory (STAI; 25) increased from before to during the COVID-19 pandemic (*t*(28) = 3.12, *p* = .004, *d* = 0.58, two-tailed). The corresponding intraclass correlation coefficient showed poor temporal stability (ICC = 0.46, 95% CI [-.01, .71]; Figure 2B). These findings suggest that even though the impact of the pandemic increased participants’ anxiety levels, curiosity traits remained the same. Given that we did not find any significant differences in curiosity traits from before to during the pandemic, all subsequent analyses use curiosity trait scores measured during the pandemic in line with the measurement of information seeking of COVID-19-related news.

**Figure 2.**
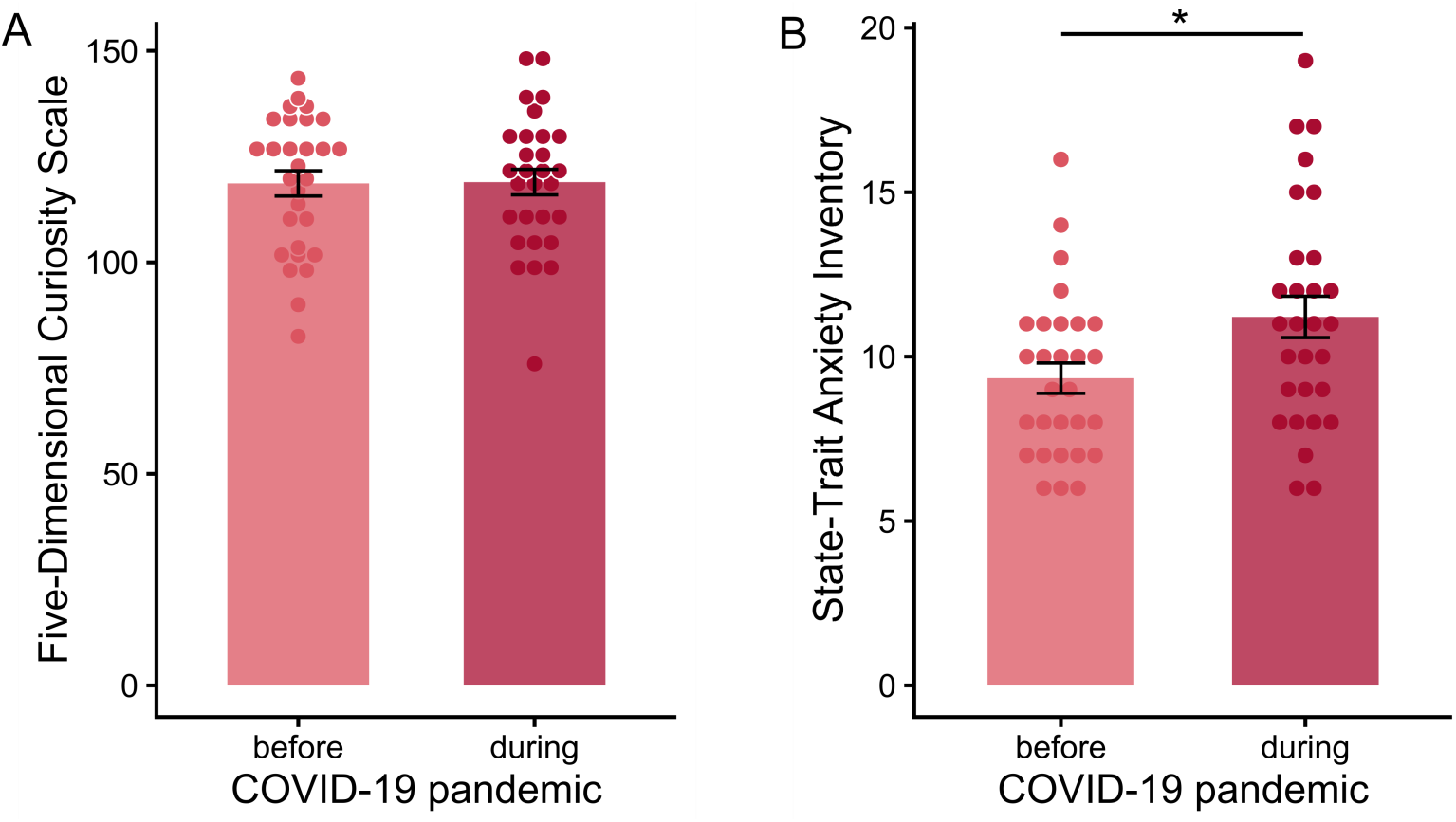
Curiosity and anxiety scores conducted before and during the COVID-19 pandemic. (A) Five-dimensional curiosity traits remained stable over time. (B) Anxiety levels increased from before to during the COVID-19 pandemic. Error bars indicate standard error of the mean.

### Does curiosity drive real-life information seeking?

In order to investigate whether curiosity positively influences real-life information seeking, we conducted one-tailed linear regressions for each of the four characteristics captured by the information seeking questionnaire. Specifically, linear regressions with five-dimensional curiosity as predictor were calculated for the frequency, detail, duration, and diversity of COVID-19-related information seeking during the first month of lockdown as dependent variables (Figure 1). Interestingly, curiosity measured by the Five-Dimensional Curiosity Scale (24) during the COVID-19 pandemic was positively associated with the frequency of information seeking during the first month of lockdown (β = 0.03, *t*(28) = 2.68, FDR-adjusted *p* = .025; Figure 3A). Thus, participants who generally experience and express high curiosity indicated by high curiosity scores informed themselves more often about COVID-19-related news per day than less curious participants. Individual differences in five-dimensional curiosity traits showed no relationship with the detail, duration, and diversity of information-seeking behaviour (all FDR-adjusted *p*-values > .142). Further examination of the five-dimensional curiosity subscales suggested that both joyous exploration (β = 0.11, *t*(28) = 2.66, FDR-adjusted *p* = .033) and deprivation sensitivity (β = 0.10, *t*(28) = 2.27, FDR-adjusted *p* = .040) drove the positive relationship between curiosity and frequency of information-seeking when adjusted for multiple comparisons correction. The remaining subscales of the five-dimensional curiosity scale were not associated with the frequency of information seeking during the first month of lockdown (all FDR-adjusted *p*-values > .092).

**Figure 3.**
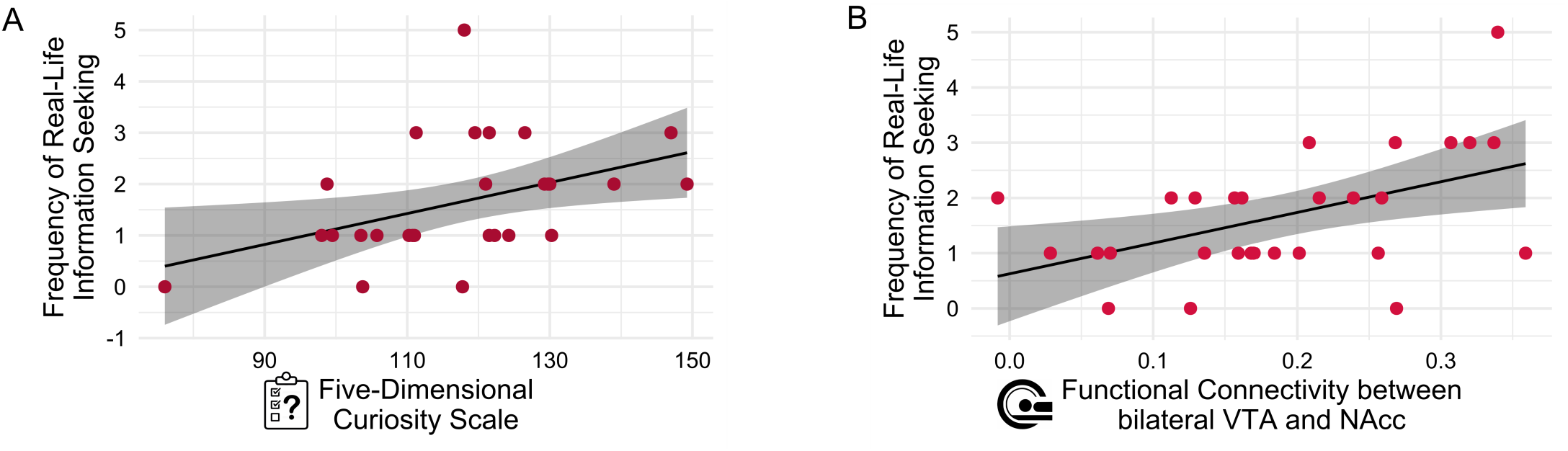
Positiv erelationship of curiosity and mesolimbic functional connectivity with the frequency of real-life information seeking. (A) Five-dimensional curiosity traits measured during the pandemic were positively associated with the frequency of COVID-19-related information seeking during the first month of lockdown. (B) Resting-state functional connectivity between bilateral VTA and NAcc was positively associated with the frequency of information-seeking behaviour.

Given that anxiety during the pandemic might have influenced information seeking as well, we conducted one-tailed linear regressions in the same way as for the assessment of curiosity. Anxiety measured during the pandemic showed no association with the frequency, detail, duration, and diversity of information-seeking behaviour (all FDR-adjusted *p*-values > .125). Furthermore, we performed a multiple regression analysis to investigate whether curiosity is still positively associated with the frequency of information seeking, when anxiety is controlled for. The multiple regression model was significant (*F*(2,26) = 3.67, *p* = .020) and explained 16.03% of variance. According to this multiple regression, solely curiosity (β = 0.03, *t*(27) = 2.70, *p* = .006) but not anxiety measured during the COVID-19 pandemic (β = −0.03, *t*(27) = −0.58, *p* = .282) was positively associated with the frequency of real-life information-seeking behaviour (Table 1).

**Table 1.** Multiple regression of curiosity and anxiety measured during the COVID-19 pandemic predicting the frequency of real-life information seeking during the first month of lockdown.

| Predictor | $\beta$ | SE | $t$ | $p$ |
| --- | --- | --- | --- | --- |
| Constant | -1.79 | 1.53 | -1.17 | .126 |
| Curiosity | 0.03 | 0.01 | 2.70 | .006** |
| Anxiety | -0.03 | 0.06 | -0.58 | .282 |

Note. \* $p < .05$ , \*\* $p < .01$ , one-tailed.

### Does mesolimbic functional connectivity drive real-life information seeking?

In order to test whether intrinsic resting-state functional connectivity (RSFC) within the mesolimbic dopaminergic circuit is positively associated with real-life information seeking, we investigated whether RSFC between bilateral VTA and NAcc predicts different characteristics of information-seeking behaviour. One-tailed linear regressions were calculated for each measure of the information seeking questionnaire. Interestingly, functional connectivity between VTA and NAcc was positively associated with the frequency of information-seeking behaviour (β = 5.50, *t*(27) = 2.75, FDR-adjusted *p* = .021; Figure 3B). This finding indicates that the stronger the functional connectivity within participants’ mesolimbic dopaminergic circuit, the more often participants informed themselves about COVID-19-related news during the first month of lockdown. RSFC between bilateral VTA and NAcc was not linked to detail, duration, and diversity of COVID-19-related information-seeking behaviour (all FDR-adjusted *p*-values > .146), suggesting a specific relationship between mesolimbic dopaminergic RSFC and the frequency of real-life information seeking.

Given that both the five-dimensional curiosity and VTA-NAcc RSFC were positively associated specifically with the frequency of COVID-19-related information seeking, we conducted a multiple regression analysis to investigate whether curiosity trait scores and bilateral VTA-NAcc RSFC were positively associated with the frequency of information seeking dependently or independently of each other (Table 2). The multiple regression model was significant (*F*(2,26) = 6.20, *p* = .003) and explained 27.07% of variance. More precisely, both curiosity (β = 0.02, *t*(27) = 1.99, *p* = .028) and RSFC between bilateral VTA and NAcc (β = 4.18, *t*(27) = 2.01, *p* = .024) were independently associated with how often participants informed themselves about COVID-19-related news per day.

**Table 2.** Multiple regression of curiosity measured during the COVID-19 pandemic and resting-state functional connectivity (RSFC) between bilateral VTA and NAcc predicting the frequency of real-life information seeking during the first month of lockdown.

| Predictor | $\beta$ | SE | $t$ | $p$ |
| --- | --- | --- | --- | --- |
| Constant | -1.89 | 1.35 | -1.40 | .087 |
| Curiosity | 0.02 | 0.01 | 1.99 | .028* |
| VTA-NAcc RSFC | 4.18 | 2.01 | 2.08 | .024* |

Note. \* $p < .05$ , \*\* $p < .01$ , one-tailed.

## DISCUSSION

The results of the present study suggest that curiosity and mesolimbic dopaminergic functional connectivity are key drivers of information seeking in real life. individual differences of both temporally stable curiosity traits and functional connectivity between bilateral VTA and NAcc were independently associated with the frequency of information seeking about COVD-19-related news. Thereby, the present results support and translate previous laboratory-based findings (1–3, 14, 17) to real-life knowledge acquisition for a specific topic that is novel and personally relevant across all information seekers. importantly, the role of curiosity in real-life information seeking held true when controlling for anxiety levels, which increased during the beginning of the COVID-19 pandemic but did not have an influence on real-life information seeking. This fits to the idea that specifically curiosity and the curiosity-promoting mesolimbic dopaminergic circuit play a crucial role in real-life information-seeking behaviour (4).

The recent COVID-19 pandemic introduced high levels of uncertainty about personally relevant consequences and information gaps about COVID-19-related information. As suggested by previous research, the detection of information gaps can elicit curiosity, leading to subsequent information seeking in order to reduce uncertainty and close information gaps (1,4–6). The present results support this idea by demonstrating that individual differences in curiosity traits, that is, the general curiosity of a person, are associated with how often participants inform themselves about COVID-19-related news. Consequently, not only temporally variable curiosity states are associated with COVID-19-related information seeking (19) but also temporally stable curiosity traits. Despite the overarching influence of the pandemic on physiological and psychological well-being (20, 21), curiosity traits, as measured with the Five-Dimensional Curiosity Scale (24), remained stable over time while anxiety levels increased from before to during the COVID-19 pandemic. This is consistent with the idea that curiosity is a rather stable personality trait that drives information-seeking behaviour in everyday life. Our finding adds to research showing that information seeking motives remain stable over time – even during such a major impact on our lives as the COVID-19 pandemic (26). In line with the finding that deprivation sensitivity – a subtype of curiosity that reflects the tendency to seek information in order to close information gaps – is associated with the creation of knowledge networks during information seeking (9), we also found a relationship between curiosity and information seeking that was driven by deprivation sensitivity. This finding further suggests that participants who experience a stronger need to close information gaps inform themselves more frequently about personally relevant information. In addition, we found that the subtype joyous exploration also predicted the frequency of real-life information seeking. Therefore, our results extend prior research, suggesting that also those who are characterised by more diverse, exploratory curiosity seek out personally relevant information more often. Even though the government regulations in response to the COVID-19 pandemic included measures of social distancing and isolation, the subscale social curiosity was not associated with real-life information seeking. Given that previous research found mixed results regarding the relationship between anxiety and information seeking, it was not clear whether curiosity alone or in addition to anxiety would be linked to real-life information seeking. Anxiety during the COVID-19 pandemic had been associated with a reduction in the willingness to wait for information (19) but also increased information-seeking behaviour (22, 23). However, anxiety was not associated with information seeking when obsessive compulsive behaviour was taken into account (23). The present results help to reconcile these inconsistent findings by providing further evidence that anxiety is not linked to real-life information seeking – not even in addition to curiosity scores. Thus, it seems to be specifically curiosity and mesolimbic dopaminergic functional connectivity that drive information seeking in real life.

In the present study, individual differences in intrinsic resting-state functional connectivity of key brain regions of the mesolimbic dopaminergic circuit, namely bilateral VTA and NAcc, were associated with the frequency of COVID-19-related information seeking. These results underline previous findings showing that curiosity and information seeking are accompanied by increased activity and stronger functional connectivity within the mesolimbic dopaminergic system (10–17). Furthermore, they suggest that even individual differences in taskin-dependent intrinsic functional connectivity within the mesolimbic circuit are associated with information seeking in real life. Participants with stronger intrinsic VTA-NAcc functional connectivity informed themselves about COVID-19-related news more often than participants with weaker mesolimbic functional connectivity. Consequently, the mesolimbic dopaminergic circuit forms not only a network that supports the drive to seek information in laboratory settings (16, 17) but is also important for information seeking in everyday life. The finding that curiosity and functional connectivity between bilateral VTA and NAcc were associated with information seeking independently of each other suggests that they are both important drivers of real-life information seeking. Although the information seeking questionnaire used in the present study assessed four characteristics of COVID-19-related information-seeking behaviour, curiosity was associated specifically with the frequency but not with the detail, duration, and diversity of information seeking. This specificity might be explained by the fact that participants had to rate their information-seeking behaviour during the first month of lockdown in hindsight. While this might have been fairly accurate for the frequency of information-seeking, participants might have misjudged the duration or diversity of their information-seeking behaviour (27), making potentially the frequency the most reliable measurement. The finding that both curiosity *and* VTA-NAcc functional connectivity were associated specifically with the frequency of information seeking further supports the validity of the frequency measure. However, it would be fruitful if future studies might use complementary, objective measures of real-life information seeking, such as mobile tracking devices.

Taken together, the present study provides evidence that curiosity and mesolimbic dopaminergic functional connectivity are key drivers of real-life information seeking, suggesting that laboratory-based findings are translatable to everyday life. Both curiosity and intrinsic functional connectivity of the mesolimbic dopaminergic circuit were positively associated with the frequency of COVID-19-related information seeking whereas anxiety levels were not linked to information seeking. Therefore, the present study paves the way for future translational studies that address important questions about the relationship between individual differences in curiosity, intrinsic functional brain connections, and real-life information seeking. Furthermore, the present findings lay the foundation for a better understanding of individual differences in cognitive and neural characteristics that shape how individuals seek out information. A better understanding of the drivers of real-life information seeking offers the possibility to potentially help individuals who exhibit excessive or detrimental information-seeking behaviour that might negatively affect their well-being or mental health.

## MATERIALS AND METHODS

### Participants

Overall, 63 healthy participants were invited to return to participate in this study after taking part in a previous study that included resting-state fMRI and curiosity measures. Of those invited, 33 responded to our request, leading to a 52.38% response rate. Four participants were excluded from statistical analyses due to excess motion artefacts during fMRI data acquisition and one of them also being an outlier in the frequency, detail, and duration of information seeking as determined by the Tukey’s method with three interquartile ranges. The final sample consisted of 29 participants (two male, mean age = 19.97 years, range = 19-21 years) with normal or corrected-to-normal vision. Participants gave written informed consent and the study was approved by the Cardiff University School of Psychology Ethics Committee.

### Experimental design

Participants took part in two experimental sessions. In the first session, which took place *before* the COVID-19 pandemic, participants carried out cognitive tasks (unrelated to this study), filled out various questionnaires, and their brain activity was measured via restingstate fMRI. This session took place up to 15 months before the start of the first official UK lockdown during the COVID-19 pandemic (i.e., before 16th March 2020). In the second session, which took place *during* the COVID-19 pandemic, participants repeated the Five-Dimensional Curiosity Scale (24; Cronbach’s α = .87) and the short version of the STAI (25; Cronbach’s α = .75) via Qualtrics (Provo, USA). Data collection during the pandemic took place between 25th June 2020 and 7th July 2020. In addition to the questionnaires, we measured the extent to which participants carried out information-seeking behaviour specific to COVID-19-pandemic content. Specifically, the information seeking questionnaire consisted of four items, which quantified the frequency, detail, duration, and diversity of COVID-19-related information seeking per day (cf. Figure 1). For example, the item probing frequency of time spent information seeking was shown as “How often did you inform yourself about COVID-19-related news? [per day]”. Participants were instructed to indicate numerically for each question the amount of information seeking they carried out. Participants were instructed as such: “We would like to know more about how you inform yourself about the COVID-19 pandemic. Please provide a numeric answer that best fits you for the requested quantity.” The internal consistency of the 4-item scale was Cronbach’s α = .76, which is deemed to be satisfactory (28).

To understand the role of curiosity traits on COVID-19-related information seeking, the Five-Dimensional Curiosity Scale (24) was administered before and during the beginning of the COVID-19 pandemic. This scale is subdivided into the subscales, joyous exploration, deprivation sensitivity, stress tolerance, thrill seeking, and social curiosity. The items were presented on a 7-point Likert scale, ranging from 1 = “does not describe me at all” to 7 = “completely describes me”, with the instruction “Below are a number of statements that describe ways in which people act and think. For each of the statements below, please indicate how well it describes you.” The Five-Dimensional Curiosity Scale was selected in order to capture a broad spectrum of personality, well-being, and social factors that can influence curiosity and information seeking. In addition to this curiosity questionnaire, the short form of the State-Trait Anxiety Inventory (STAI; 25) was conducted before and during the COVID-19 pandemic. Participants were given the prompt “A number of statements which people have used to describe themselves are given below. Read each statement and then select the most appropriate number to the right of the statement to indicate how you feel right now, at this moment. There are no right or wrong answers. Do not spend too much time on any one statement but give the answer which seems to describe your present feelings best.” The scales used were 1 = “not at all”, 2 = “somewhat”, 3 = “moderately”, and 4 = “very much”. Items included in this scale included “I feel calm”, “I feel content”, and “I am worried”.

### FMRI acquisition

Imaging data were obtained at CUBRIC, Cardiff University, using a Siemens Magnetom Prisma 3T MRI scanner with a 32-channel head coil. High-resolution T_1_-weighted structural images were obtained using an MPRAGE sequence (TR = 2500 ms, TE = 3.06 ms, flip angle = 9°, FoV = 256 mm^2^, voxel-size = 1 mm^3^, slice thickness = 1 mm, 224 sagittal slices, bandwidth = 230 Hz/pixel; acquisition time = 7.36 min). During the structural scan, participants watched a film in order to help reduce movement, boredom, and nervousness. For resting-state fMRI, fifty transversal slices were acquired by using an echoplanar imaging (EPI) sequence (TR = 3000 ms, TE = 30 ms, flip angle 89°, FoV = 192 mm^2^, voxel-size = 2 mm^3^, slice thickness = 2 mm, bandwidth = 2170 Hz/pixel; acquisition time = 10.11 min). A black fixation cross centred on a grey background was presented during resting-state fMRI acquisition. Participants were instructed to keep their eyes open, fixate on the cross, and try to the best of their ability to keep their minds clear. They were told not to linger on things that came to their mind (29).

### Resting-state functional connectivity pre-processing and analysis

Resting-state fMRI data were pre-processed using the CONN toolbox (version 18b; 30), in conjunction with the Statistical Parametric Mapping (SPM12) modules (Wellcome Trust Centre for Neuroimaging, London) executed in MATLAB (version 2015). In a first step, functional scans were realigned and resampled to a reference structural image using SPM12 (31) and slice-time corrected (32), adjusting for differences in acquisition times between the inter-leaved scans. The Artefact Detection Tool (ART) was used to flag potential outliers with a framewise displacement > 0.5 mm and global BOLD signal change exceeding three standard deviations of subject-specific means. Then, structural and functional images were normalised into MNI space and segmented (33), with 2 mm isotropic voxels for functional and 1 mm isotropic voxels for structural images. Functional imaging was spatially smoothed using 6 mm full-width-half maximum (FWHM) Gaussian Kernel. Next, images were denoised using CONN’s anatomical component noise correction procedure (34, 35). Twelve noise components (three translation, three rotation parameters, and their respective first-order derivatives) were identified (36) to reduce motion variability in the BOLD signal. Outlier scans identified in ART were scrubbed during this step. To remove slow trends in the signal and initial magnetization transients from the BOLD signal, a linear detrending was used. Finally, the standard band-pass filter between 0.008 Hz and 0.09 Hz was used. In addition, data from participants with over 15% of invalid scans as identified by ART were removed from analysis. According to this exclusion criterion, data from four participants were removed.

In order to investigate intrinsic functional connectivity between VTA and NAcc, bilateral masks of both brain regions were used as regions of interest (ROIs). The bilateral mask of the VTA was taken from Murty et al. (37). Left and right NAcc masks were taken from the Harvard-Oxford cortical atlas (38) and combined to a bilateral NAcc mask in FSL (39). The BOLD time series of each ROI was computed by averaging the voxel time series across all voxels within the ROI. Each participant’s intrinsic mesolimbic functional connectivity was computed as Fisher’s z-transformed bivariate Pearson correlation coefficient between the VTA and NAcc BOLD time series.

### Data analyses

Changes in curiosity and anxiety from before to during the COVID-19 pandemic were investigated using repeated-measures *t*-tests. To further understand whether the measurements are temporally stable, intraclass correlations were carried out with the ranges < 0.5, 0.5-0.75, and 0.75-0.9 indicating poor, moderate, and good temporal reliabilities, respectively (40). To understand to what extent curiosity influences characteristics of COVID-19-related information seeking, we carried out linear regressions. In addition, to control that this was not driven by underlying anxiety, we carried out a multiple regression. Finally, as we expected that these factors would contribute to real-life information seeking, we examined a potential positive relationship between curiosity and resting-state functional connectivity between bilateral VTA and NAcc. As such, a multiple regression was carried out to understand whether these predictors were dependent or independent of each other. For all analyses, the significance level was set to α = .05 and if not indicated differently, one-tailed results are reported given our directed hypotheses. In order to correct for multiple comparisons, the false discovery rate (FDR) method was applied and adjusted *p*-values were reported (41). In order to avoid biases from outliers for all statistical analyses, outliers were detected with the Tukey’s method using three interquartile ranges (42). One participant was classified as an outlier for the frequency, detail, and duration of information seeking and was removed from analyses also due to excess motion artefacts.

## ACKNOWLEDGEMENTS

The authors thank Bonni Crawford and Alisa Priemysheva for their help with the study design. The study was supported by a Wellcome Trust and Royal Society Sir Henry Dale Fellowship (211201/Z/18/Z) awarded to M.J.G. K.C.J.E. received funding by Wellcome Trust (AC1710IF09) and the German Research Foundation (442588275). D.F.M.M.P., A.V., and V.D. were supported by PhD studentships funded by the Cardiff University School of Psychology. For the purpose of Open Access, the authors have applied a CC-BY public copyright license to any Author Accepted Manuscript version arising from this submission.

## REFERENCES

1. J. Gottlieb, P.-Y. Oudeyer, M. Lopes, A. Baranes, Information-seeking, curiosity, and attention: Computational and neural mechanisms. Trends Cogn. Sci. 17, 585–593 (2013).

2. C. Kidd, B. Y. Hayden, The Psychology and Neuroscience of Curiosity. Neuron 88, 449–460 (2015).

3. L. L. F. van Lieshout, F. P. de Lange, R. Cools, Why so curious? Quantifying mechanisms of information seeking. Current Opinion in Behavioral Sciences 35, 112–117 (2020).

4. M. J. Gruber, C. Ranganath, How Curiosity Enhances Hippocampus-Dependent Memory: The Prediction, Appraisal, Curiosity, and Exploration (PACE) Framework. Trends Cogn. Sci. 23, 1014–1025 (2019).

5. G. Loewenstein, The psychology of curiosity: A review and reinterpretation. Psychological Bulletin 116, 75–98 (1994).

6. L. van Lieshout, I. Traast, F. de Lange, R. Cools, Curiosity or savouring? Information seeking is modulated by both uncertainty and valence. PLOS ONE 16, e0257011 (2021).

7. P. J. Silvia, What Is Interesting? Exploring the Appraisal Structure of Interest. Emotion 5, 89–102 (2005).

8. M. K. Noordewier, E. van Dijk, Interest in Complex Novelty. Basic and Applied Social Psychology 38, 98–110 (2016).

9. D. M. Lydon-Staley, D. Zhou, A. S. Blevins, P. Zurn, D. S. Bassett, Hunters, busybodies and the knowledge network building associated with deprivation curiosity. Nat Hum Behav 5, 327–336 (2021).

10. M. J. Gruber, B. D. Gelman, C. Ranganath, States of curiosity modulate hippocampus-dependent learning via the dopaminergic circuit. Neuron 84, 486–496 (2014).

11. M. J. Kang, et al., The wick in the candle of learning: epistemic curiosity activates reward circuitry and enhances memory. Psychol. Sci. 20, 963–973 (2009).

12. R. Ligneul, M. Mermillod, T. Morisseau, From relief to surprise: Dual control of epistemic curiosity in the human brain. Neuroimage 181, 490–500 (2018).

13. S. Oosterwijk, L. Snoek, J. Tekoppele, L. H. Engelbert, H. S. Scholte, Choosing to view morbid information involves reward circuitry. Sci. Rep. 10, 15291 (2020).

14. J. K. L. Lau, H. Ozono, K. Kuratomi, A. Komiya, K. Murayama, Shared striatal activity in decisions to satisfy curiosity and hunger at the risk of electric shocks. Nat Hum Behav 4, 531–543 (2020).

15. J.-H. Poh, et al., Tuned to Learn: An anticipatory hippocampal convergence state conducive to memory formation revealed during midbrain activation. bioRxiv [Preprint] (2021) https:/doi.org/10.1101/2021.07.15.452391

16. L. FitzGibbon, J. K. L. Lau, K. Murayama, The seductive lure of curiosity: Information as a motivationally salient reward. Current Opinion in Behavioral Sciences 35, 21 – 27 (2020).

17. C. J. Charpentier, E. S. Bromberg-Martin, T. Sharot, Valuation of knowledge and ignorance in mesolimbic reward circuitry. Proc. Natl. Acad. Sci. U. S. A. 115, E7255–E7264 (2018).

18. I. Kahn, D. Shohamy, Intrinsic connectivity between the hippocampus, nucleus accumbens, and ventral tegmental area in humans. Hippocampus 23, 187–192 (2013).

19. Y. Abir, et al., Rational Curiosity and Information-Seeking in the COVID-19 Pandemic. PsyArXiv [Preprint] (2020) https:/doi.org/10.31234/osf.io/hcta4

20. M. J. Savage, et al., Mental health and movement behaviour during the COVID-19 pandemic in UK university students: Prospective cohort study. Mental Health and Physical Activity 19, 100357 (2020).

21. C. Scrivner, J. A. Johnson, J. Kjeldgaard-Christiansen, M. Clasen, Pandemic practice: Horror fans and morbidly curious individuals are more psychologically resilient during the COVID-19 pandemic. Pers. Individ. Dif. 168, 110397 (2021).

22. C. J. Charpentier, et al., Anxiety selectively increases information-seeking in response to large changes. PsyArXiv [Preprint] (2021) https:/doi.org/10.31234/osf.io/n2puk

23. A. M. Loosen, V. Skvortsova, T. U. Hauser, Obsessive-compulsive symptoms and information seeking during the Covid-19 pandemic. Transl. Psychiatry 11, 309 (2021).

24. T. B. Kashdan, et al., The five-dimensional curiosity scale: Capturing the bandwidth of curiosity and identifying four unique subgroups of curious people. J. Res. Pers. 73, 130–149 (2018).

25. T. M. Marteau, H. Bekker, The development of a six-item short-form of the state scale of the Spielberger State-Trait Anxiety Inventory (STAI). Br. J. Clin. Psychol. 31, 301–306 (1992).

26. C. A. Kelly, T. Sharot, Individual differences in information-seeking. Nat. Commun. 12, 7062 (2021).

27. D. A. Parry, et al., A systematic review and meta-analysis of discrepancies between logged and self-reported digital media use. Nat Hum Behav 5, 1535–1547 (2021).

28. M. Tavakol, R. Dennick, Making sense of Cronbach’s alpha. J. Int. Assoc. Med. Sci. Educ. 2, 53–55 (2011).

29. B. B. Biswal, J. Van Kylen, J. S. Hyde, Simultaneous assessment of flow and BOLD signals in resting-state functional connectivity maps. NMR Biomed. 10, 165–170 (1997).

30. S. Whitfield-Gabrieli, A. Nieto-Castanon, Conn: a functional connectivity toolbox for correlated and anticorrelated brain networks. Brain Connect. 2, 125–141 (2012).

31. J. L. Andersson, C. Hutton, J. Ashburner, R. Turner, K. Friston, Modeling geometric deformations in EPI time series. Neuroimage 13, 903–919 (2001).

32. R. N. A. Henson, C. Buechel, O. Josephs, K. J. Friston, The slice-timing problem in event-related fMRI. Neuroimage 9, 125 (1999).

33. J. Ashburner, K. J. Friston, Unified segmentation. Neuroimage 26, 839–851 (2005).

34. Y. Behzadi, K. Restom, J. Liau, T. T. Liu, A component based noise correction method (CompCor) for BOLD and perfusion based fMRI. Neuroimage 37, 90–101 (2007).

35. X. J. Chai, A. N. Castañón, D. Öngür, S. Whitfield-Gabrieli, Anticorrelations in resting state networks without global signal regression. Neuroimage 59, 1420–1428 (2012).

36. K. J. Friston, S. Williams, R. Howard, R. S. J. Frackowiak, R. Turner, Movement-related effects in fMRI time-series. Magn. Reson. Med. 35, 346–355 (1996).

37. V. P. Murty, et al., Resting state networks distinguish human ventral tegmental area from substantia nigra. Neuroimage 100, 580–589 (2014).

38. G. Grabner, et al., Symmetric atlasing and model based segmentation: an application to the hippocampus in older adults. Med. Image Comput. Comput. Assist. Interv. 9, 58–66 (2006).

39. M. Jenkinson, C. F. Beckmann, T. E. J. Behrens, M. W. Woolrich, S. M. Smith, FSL. Neuroimage 62, 782–790 (2012).

40. T. K. Koo, M. Y. Li, A Guideline of Selecting and Reporting Intraclass Correlation Coefficients for Reliability Research. J. Chiropr. Med. 15, 155–163 (2016).

41. Y. Benjamini, Y. Hochberg, Controlling the false discovery rate: A practical and powerful approach to multiple testing. J. R. Stat. Soc. 57, 289–300 (1995).

42. J. W. Tukey, Exploratory Data Analysis (Addison-Wesley, 1977).

